# Poly(A)-binding protein is an ataxin-2 chaperone that emulsifies biomolecular condensates

**DOI:** 10.1101/2021.08.23.457426

**Authors:** Steven Boeynaems, Yanniv Dorone, Anca Marian, Victoria Shabardina, Guozhong Huang, Garam Kim, Anushka Sanyal, Nesli-Ece Şen, Roberto Docampo, Iñaki Ruiz-Trillo, Keren Lasker, Georg Auburger, Edor Kabashi, Aaron D. Gitler

**Affiliations:** Department of Genetics, Stanford University School of Medicine, Stanford, California 94305, USA; Department of Plant Biology, Carnegie Institution for Science, Stanford, California 94305, USA; Department of Biology, Stanford University, Stanford, CA 94305, USA; Imagine Institute, Institut National de la Santé et de la Recherche Médicale (INSERM) Unité 1163, Paris Descartes Université, 75015 Paris, France; Institut de Biologia Evolutiva (CSIC-Universitat Pompeu Fabra), Passeig Marítim de la Barceloneta 37-49, 08003 Barcelona, Catalonia, Spain; Department of Cellular Biology and Center for Tropical and Emerging Global Diseases, University of Georgia, Athens, Georgia 30602, USA; Experimental Neurology, Goethe-University Hospital, 60590 Frankfurt, Germany; ICREA, Passeig Lluís Companys 23, 08010 Barcelona, Catalonia, Spain; The Scripps Research Institute, La Jolla, CA 92037

**Keywords:** Phase separation, short linear motif, polyQ, protein aggregation, prion-like, stress granules, frontotemporal dementia, amyotrophic lateral sclerosis, spinocerebellar ataxia

## Abstract

Biomolecular condensation underlies the biogenesis of an expanding array of membraneless assemblies, including stress granules (SGs) which form under a variety of cellular stresses. Advances have been made in understanding the molecular grammar that dictates the behavior of a few key scaffold proteins that make up these phases but how the partitioning of hundreds of other SG proteins is regulated remains largely unresolved. While investigating the rules that govern the condensation of ataxin-2, a SG protein implicated in neurodegenerative disease, we unexpectedly identified a short 14aa sequence that acts as an ataxin-2 condensation switch and is conserved across the eukaryote lineage. We identify poly(A)-binding proteins as unconventional RNA-dependent chaperones that control this regulatory switch. Our results uncover a hierarchy of *cis* and *trans* interactions that fine-tune ataxin-2 condensation and reveal a new molecular function for ancient poly(A)-binding proteins as emulsifiers of biomolecular condensate proteins. These findings may inspire novel approaches to therapeutically target aberrant phases in disease.

## INTRODUCTION

Stress granules (SGs) are cytoplasmic ribonucleoprotein assemblies that form when cells are exposed to stress (e.g., heat, chemical, osmotic stress) (Wolozin and Ivanov, 2019). SGs were initially thought to function in translational regulation, but emerging evidence indicates that they may serve a protective role in the cell by preventing the irreversible aggregation of proteins and even mRNA (Guillen-Boixet et al., 2020; Mann et al., 2019; McGurk et al., 2018). Hundreds of proteins and thousands of mRNAs have been found to localize to SGs (Jain et al., 2016; Khong et al., 2017; Markmiller et al., 2018; Marmor-Kollet et al., 2020; Namkoong et al., 2018). Many of these SG-localized proteins harbor “sticky” and aggregation-prone domains (Boeynaems et al., 2017a; King et al., 2012; Kuechler et al., 2020), and the ability of these proteins to interact with RNA might help protect them from aggregation (Maharana et al., 2018; Mann et al., 2019). Given their aggregation-prone nature, several SG proteins have been implicated in human protein aggregation diseases. For instance, missense mutations in RNA-binding proteins TDP-43, FUS, EWS1, TAF15, and other heterogeneous nuclear ribonucleoproteins (hnRNPs) are linked to hereditary forms of degenerative diseases amyotrophic lateral sclerosis (ALS), frontotemporal dementia (FTD), and myopathies (Li et al., 2013; Ramaswami et al., 2013). Additionally, loss-of-function mutations have been found in proteins that regulate SG clearance (Buchan et al., 2013; Monahan et al., 2016), indicating that while SGs might be initially protective, their misregulation may underlie eventual pathological aggregation (Li et al., 2013; Ramaswami et al., 2013).

An emerging view is that SGs and other so-called biomolecular condensates form via phase separation (Boeynaems et al., 2018; Shin and Brangwynne, 2017). The set of rules dictating this behavior remains unresolved but for RNA-centric condensates (including SGs), a picture emerges where condensation is driven by a complex interplay of strong specific and weak promiscuous interactions. Recent work has uncovered a hierarchy of such interactions between oligomerization, RNA-binding, and disordered domains, and elucidated how this these rules govern the phase separation of key SG scaffold proteins such as G3BP1 (Guillen-Boixet et al., 2020; Sanders et al., 2020; Yang et al., 2020). Additionally, the disordered sticky prion-like domains of several ALS-related RNA-binding proteins are involved in the liquid-like behavior of these proteins (Lin et al., 2015; Molliex et al., 2015; Patel et al., 2015). Whether these same simple rules apply to other SG proteins remains unknown. Defining these principles will be crucial for devising novel ways to therapeutically modulate phase separation and aggregation of these disease-linked proteins.

Ataxin-2 (ATXN2), another SG-associated protein, has been implicated in two neurodegenerative diseases – spinocerebellar ataxia-2 (SCA2) and ALS. ATXN2 harbors a polyglutamine (polyQ) domain. Long expansions (>34 repeats) of this polyQ domain cause SCA2 (Imbert et al., 1996; Pulst et al., 1996; Sanpei et al., 1996), whereas intermediate-length repeat expansions (22-34 repeats) are associated with ALS (Elden et al., 2010). Notably, lowering levels of ATXN2 is sufficient to mitigate neurodegeneration and extend survival in multiple model systems (Becker et al., 2017; Elden et al., 2010; Scoles et al., 2017). Mechanistically, ATXN2 localizes to SGs and promotes TDP-43 mislocalization (Becker et al., 2017). The protein has also been implicated in dendritogenesis and memory formation in flies, potentially by regulating mRNA stability and transport (Bakthavachalu et al., 2018; Singh et al., 2021). These functions are dependent on the ability of ATXN2 to condense into RNA granules via its intrinsically disordered regions (IDRs). Additionally, ATXN2 has been implicated in the regulation of circadian rhythm in flies (Lim and Allada, 2013; Zhang et al., 2013) and mice (Pfeffer et al., 2017), and perturbed sleeping patterns have been observed in both SCA and ALS cases (Boentert, 2020; Huebra et al., 2019). Thus, defining the mechanisms connecting ATXN2 to condensate form and function will provide insight into this important disease gene and could suggest additional ways to safely target this function therapeutically (Auburger et al., 2017).

Here we set out to define the molecular determinants regulating ATXN2’s phase separation behavior. ATXN2 is a 140kDa intrinsically disordered RNA-binding protein, but contrary to expectation we found that its SG targeting is exclusively mediated by a short linear motif (SLiM) that confers interaction with cytoplasmic poly(A)-binding proteins (PABPCs). We provide evidence that PABPC acts as an ATXN2 chaperone by preventing its spontaneous condensation under non-stress conditions. Through evolutionary analysis, we uncover the hierarchy of *cis* and *trans* molecular interactions that regulate ATXN2 condensation. The identification of PABPC as an RNA-dependent chaperone provides an unexpected novel function for this essential protein. These findings have direct implications for our understanding of the complex regulatory networks governing protein condensation, and may inspire novel therapeutic approaches to target and modulate this behavior in human disease.

## RESULTS

### The PAM2 motif acts as a switch that regulates ATXN2 condensation in time and space

Hundreds of proteins condense into SGs upon cellular stress (Fig. 1A), including ATXN2. But the functional contributions of each of these proteins to condensation remain largely unknown. Some of these have been shown to be essential for SG formation (Kedersha et al., 2016; Yang et al., 2020), so-called scaffold proteins, whereas others are called client proteins and partition into these condensates but are not required for their formation. To test if ATXN2 functions as a SG scaffold or client, we generated an ATXN2 knock-out HeLa cell line. These cells were still able to form SGs upon treatment with arsenite, indicating that ATXN2 acts as a client protein (Fig. 1B-C), consistent with previous observations from knock-down studies (Becker et al., 2017; Lastres-Becker et al., 2016). To define the molecular features that drive ATXN2 to partition into SGs, we performed systematic domain deletions and evaluated their effect on SG partitioning in U2OS cells (Fig. 1D). ATXN2 consists of two RNA-binding domains (Lsm and LsmAD), three intrinsically disordered domains (IDR1, 2 and 3), a polyQ repeat, and a PAM2 short linear motif (SLiM), which mediates ATXN2’s interaction with cytoplasmic poly(A)-binding protein (PABPC) (Jimenez-Lopez and Guzman, 2014; Kozlov et al., 2010; Xie et al., 2014). Surprisingly, whereas RNA-binding and disordered domains have been extensively implicated in protein phase separation (Lin et al., 2015; Molliex et al., 2015; Patel et al., 2015), these domains were dispensable for SG targeting and mixing of ATXN2 into the condensed phase (measured as heterogeneity, see STAR Methods) (Fig. 1E-F). However, deleting the PAM2 SLiM, a 14 amino acid sequence that makes up only ∼1% of the ATXN2 sequence, prevented proper mixing of ATXN2 into SGs. Instead, ΔPAM2 ATXN2 formed small condensates that dotted the surface of SGs (Fig. 1E-F). Because ΔPAM2 ATXN2 demixed from SGs, we tested if its condensation was stress-dependent. Expressing this mutant in non-stressed cells resulted in spontaneous condensation under standard conditions (Fig. 1G). Since high levels of wildtype ATXN2 expression can lead to aggregation, we confirmed that the ΔPAM2 ATXN2 protein was expressed at comparable levels to the WT protein (Fig. 1H). We performed fluorescence recovery after photobleaching (FRAP) experiments and found that both spontaneous wildtype and mutant condensates have solid-like properties (Fig. 1I). Thus, our findings indicate that the PAM2 motif acts as a switch for the spatiotemporal regulation of ATXN2’s phase separation behavior – PAM2 is required for (1) proper targeting and mixing of ATXN2 into SGs under times of stress, and (2) preventing its spontaneous condensation into solid-like gels under normal conditions.

**Figure 1:**
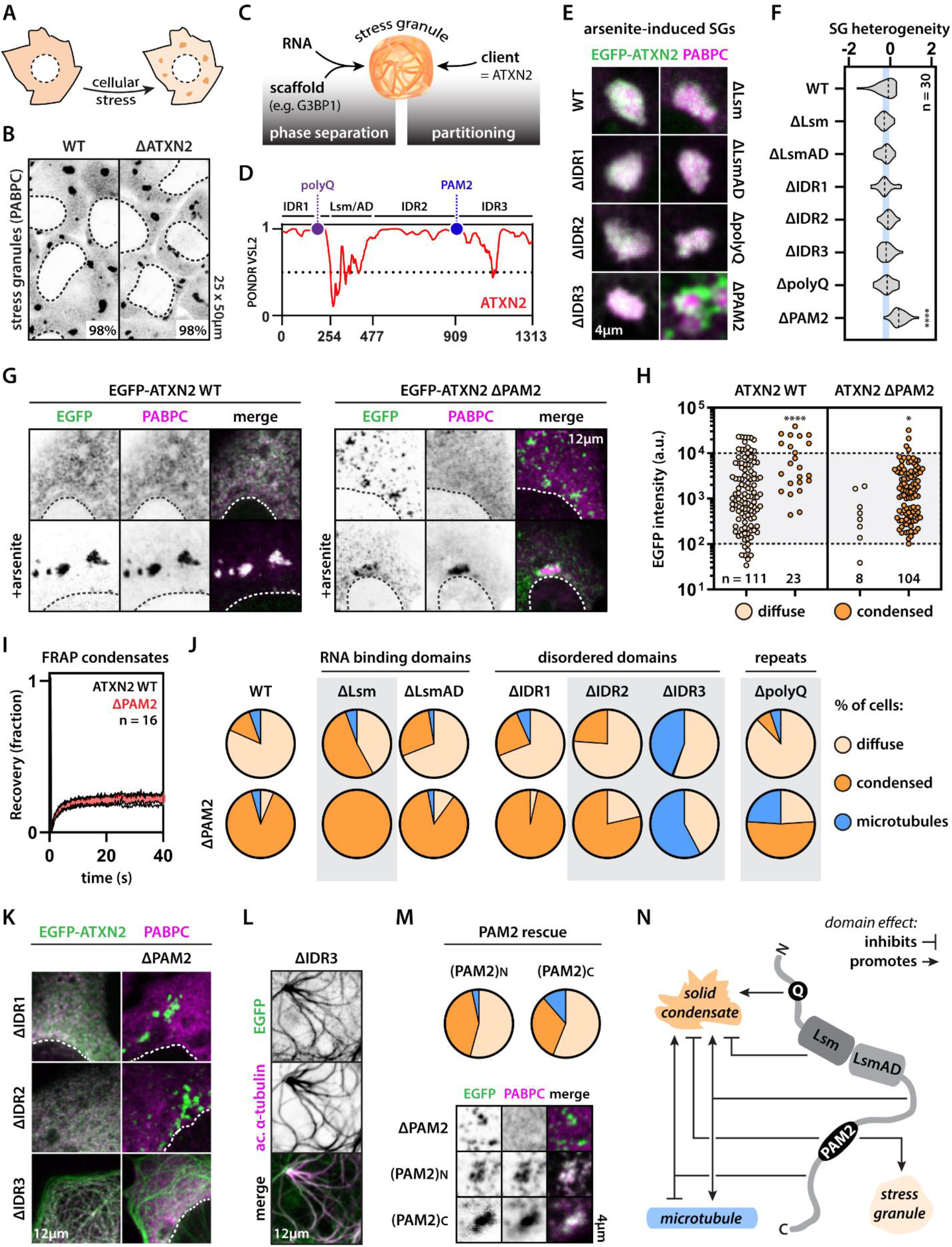
The PAM2 switch regulates ATXN2’s behavior in stress and non-stress conditions. (A) Stress granules condense upon cellular stress. (B) ATXN2 is dispensable for arsenite-induced stress granule assembly. Average percentage of cells with stress granules is shown. n = 3 experiments with a total of [255-285] cells. Dashed lines highlight nuclei. EGFP is shown in inverse gray scale. (C) Phase separation of scaffold proteins and RNA drives stress granule assembly, with subsequent recruitment of non-essential client proteins, such as ATXN2. (D) Domain structure and disorder prediction of ATXN2. PONDR score: 1 = disordered, 0 = folded. (E) Deletion of PAM2 prevents homogeneous partitioning of ATXN2 in stress granules. (F) Quantification of the heterogeneity of ATXN2 deletion mutant distribution within the stress granule compartment (see M&M). One-way ANOVA. n = 30 stress granules. (G) PAM2 deletion drives spontaneous condensation of ATXN2 into small granules under non-stress conditions. (H) Wildtype ATXN2 can spontaneously condense upon overexpression. ΔPAM2 ATXN2 condensation is not an overexpression artefact. Scatterplots show cells with diffuse or condensed ATXN2 localization. Cells combined from 3 experiments. Mann-Whitney. (I) ATXN2 is largely immobile in both wildtype and ΔPAM2 spontaneous condensates. n = 16 condensates. (J) Differential contribution of ATXN2 domains to its phase behavior. Grey boxes highlight domain deletions that strongly affect ATXN2 behavior. Cells combined from 3 experiments. n = [96-155] cells (see also Fig. S1). (K) Example pictures of IDR deletion and IDR-PAM2 double deletion mutants. (L) Example picture showing microtubule localization (acetylated tubulin antibody) of the IDR3 deletion mutant. (M) PAM2 position along the ATXN2 sequence is important for preventing spontaneous condensation. n = [123-179] cells (see also Fig. S1). (N) Scheme highlighting the complex interactions between different domains on ATXN2 behavior. Panel (B) shows HeLa cells. All other panels show U2OS cells. * p-value < 0.05, **** p-value < 0.0001.

### Complex interactions between ATXN2 domains dictate its biophysical behavior

Our initial set of domain deletion mutants identified ATXN2’s PAM2 motif as the key regulator of its phase separation behavior. Next, we sought to define how this regulatory switch interacts with other ATXN2 domains (Fig. 1J, Fig S1A). Deletion of the main RNA-binding domain LSM potentiated the spontaneous condensation of both the single (ΔLSM) and double (ΔPAM2-ΔLSM) mutants (Fig. 1J). This observation is in line with recent work on other prion-like proteins showing that RNA binding can prevent their spontaneous condensation (Maharana et al., 2018; Mann et al., 2019; Schmidt and Rohatgi, 2016; Yu et al., 2021). Conversely, deletion of the polyQ repeat, an established driver of aggregation (Lieberman et al., 2019), decreased condensation of ΔPAM2 ATXN2. Interestingly, a significant population of cells exhibited localization of ATXN2 to microtubules (Fig. 1J). We next tested the impact of each of the three IDRs. Deleting IDR1 (ΔIDR1) did not have a strong impact on ATXN2 condensation (despite the fact that this IDR contains the polyQ repeat), while ΔIDR2 and ΔIDR3 had opposing effects on ATXN2’s behavior (Fig. 1J-K). Deletion of IDR2 modestly reduced condensation of the ΔPAM2 mutant, but more importantly, abolished any microtubule localization of both the single and double mutant. Lastly, deletion of IDR3 completely prevented spontaneous condensation of ATXN2, and conferred microtubule localization in about half of the cells regardless of the presence of the PAM2 motif (Fig. 1J-L).

Since IDR2 and IDR3 had opposing effects on promoting condensation versus microtubule targeting, we wondered whether the precise localization of the PAM2 switch in between these IDRs was of functional importance. To test this, we generated add-back mutants where we reintroduced the PAM2 motif to the N- or C-terminus of a ΔPAM2 mutant. Even though both of these mutants were able to bind PABPC, they only partially rescued the spontaneous condensation phenotype (Fig. 1M, Fig. S1B), suggesting that the precise interplay between different ATXN2 domains is important for its proper regulation (Fig. 1N).

### The PAM2 switch is conserved across eukaryotes

The divergent behavior of IDR2 and IDR3, combined with our add-back experiment, suggested that the location of the PAM2 mutant is important. Since ATXN2 is a pan-eukaryote protein, we performed an evolutionary analysis. The PAM2 motif is always centered between IDR2 and IDR3 (Fig. S2A), in contrast to the polyQ repeat, which can be found at different locations in different lineages or is sometimes even absent (Jimenez-Lopez and Guzman, 2014). Given the striking conservation of the PAM2 motif, we wondered whether its role as a regulatory switch would also be conserved (Fig. 2A). We tested this in two protists, where we identified and synthesized their respective ATXN2 orthologs and generated ΔPAM2 mutants. First, *Capsaspora owczarzaki* is a close unicellular relative of animals and a model for the study of the origin of multicellularity (Ferrer-Bonet and Ruiz-Trillo, 2017). Wildtype *Co*ATXN2 (CAOG_07908) spontaneously formed several small granules, whereas PAM deletion resulted in the formation of one single large granule (Fig. 2B, Fig. S2B-D). Second, *Trypanosoma brucei*, the parasite causing sleeping sickness, is a member of the basal eukaryote lineage *Discoba*. Wildtype *Tb*ATXN2 (Tb927.8.4540) localized diffusely to the cytoplasm and PAM2 deletion resulted in the appearance of spontaneous condensates (Fig. 2C, Fig. S2E). To extend this analysis to other multicellular eukaryotes, we performed the same experiment in the model plant *Arabidopsis thaliana*. We generated stable transgenic lines carrying GFP-tagged wildtype and ΔPAM2 *At*ATXN2 (CID4) (Fig. 2D). By assaying cotyledon (embryonic leaf) and root tissue in *Arabidopsis* seedlings, we found that the PAM2 motif prevents spontaneous condensation *in vivo*. To evaluate whether the PAM2 motif was also required for SG targeting, we transiently co-infected tobacco leaves with *At*ATXN2-GFP and PAB2-RFP, which is the *Arabidopsis* ortholog of PABPC (Fig. 2E). Identical to our observations in human cells, wildtype *At*ATXN2 partitioned with PABPC-positive SGs, whereas ΔPAM2 condensates did not mix with SGs. Together, these data indicate that the PAM2 switch modulates ATXN2 condensation across eukaryotes, providing evidence for the biological importance of this ancient regulatory element.

**Figure 2:**
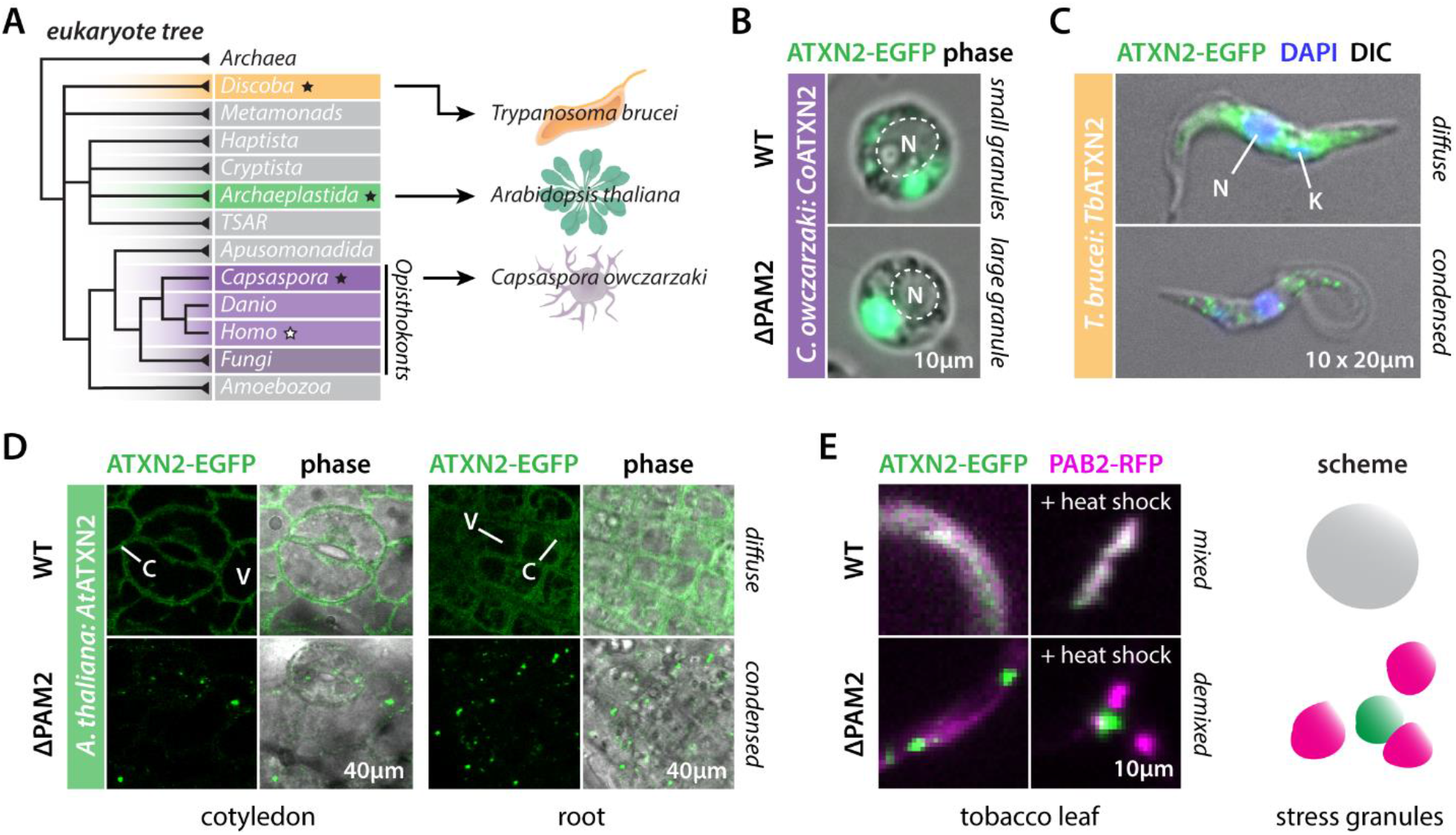
The PAM2 switch is functionally conserved across eukaryotes. (A) Eukaryote tree highlighting different clades and model organisms. Archaea are the outgroup. Black stars indicate organisms compared in this figure. (B) PAM2 deletion results in the formation of a single large *Co*ATXN2 granule upon expression in *C. owczarzaki*, opposed to multiple small wildtype granules. N denotes nucleus (see also Fig. S2B-D). (C) *Tb*ATXN2 localizes diffusely in trypanosomes, whereas the ΔPAM2 mutant spontaneously condenses (see also Fig. S2E). N denotes nucleus, K denotes kinetoplast. (D) *At*ATXN2 localizes diffusely to the cytoplasm *in vivo*, whereas ΔPAM2 *At*ATXN2 spontaneously condenses. Data are shown for the cotyledon (embryonic leaf) and root of 3-day old *A. thaliana* seedling. V denotes vacuole, C denotes cytoplasm. (E) Expression of wildtype and ΔPAM2 *At*ATXN2 in tobacco leaves recapitulates phenotypes from *A. thaliana* seedlings. PAB2 (*At*PABPC) localizes diffusely to the cytoplasm, and targets stress granules upon heat shock (30 min @ 37°C). Wildtype *At*ATXN2 partitions into PAB2 stress granules, whereas ΔPAM2 *At*ATXN2 remains demixed.

### A balance in IDR interactions controls ATXN2’s behavior

To define how the PAM2 motif counteracts spontaneous condensation, we next investigated the divergent effects of IDR2 and IDR3 deletion. IDRs are often poorly conserved at the sequence level (Brown et al., 2010), but they do tend to display conservation in amino acid composition (Zarin et al., 2019). By comparing ATXN2 IDRs spanning 1.7 billion years of eukaryote evolution, we observed that IDR2 and IDR3 have conserved differences in their composition (Fig. 3A); IDR2 is enriched in basic amino acids whereas IDR3 is enriched in aromatic residues (Fig. 3B-C). We and others have previously reported on the role of basic and aromatic residues in the condensation of SG proteins through the formation of cation-pi and pi-pi interactions (Bogaert et al., 2018; Qamar et al., 2018; Wang et al., 2018). We synthesized peptides derived from IDR2 and IDR3 and observed that they spontaneously condense when mixed together (Fig. 3D). This suggests that their interaction can indeed drive ATXN2 condensation. Thus, we propose that by physically separating these “sticky” amino acids across two IDRs, and by having the PAM2 motif separating them, ATXN2 can regulate the interaction strength between these IDRs. In this model (Fig. 3G), PABPC provides steric hindrance to the IDR2-IDR3 interaction, which is in line with the spontaneous condensation of the ΔPAM2 mutant. This model predicts that removing the physical separation between basic and aromatic residues should circumvent the effects of PABPC binding. Indeed, when scrambling the sequence of IDR2 and IDR3, while keeping the PAM2 motif intact, we observe increased condensation for this ATXN2 variant (Fig. S1B).

**Figure 3:**
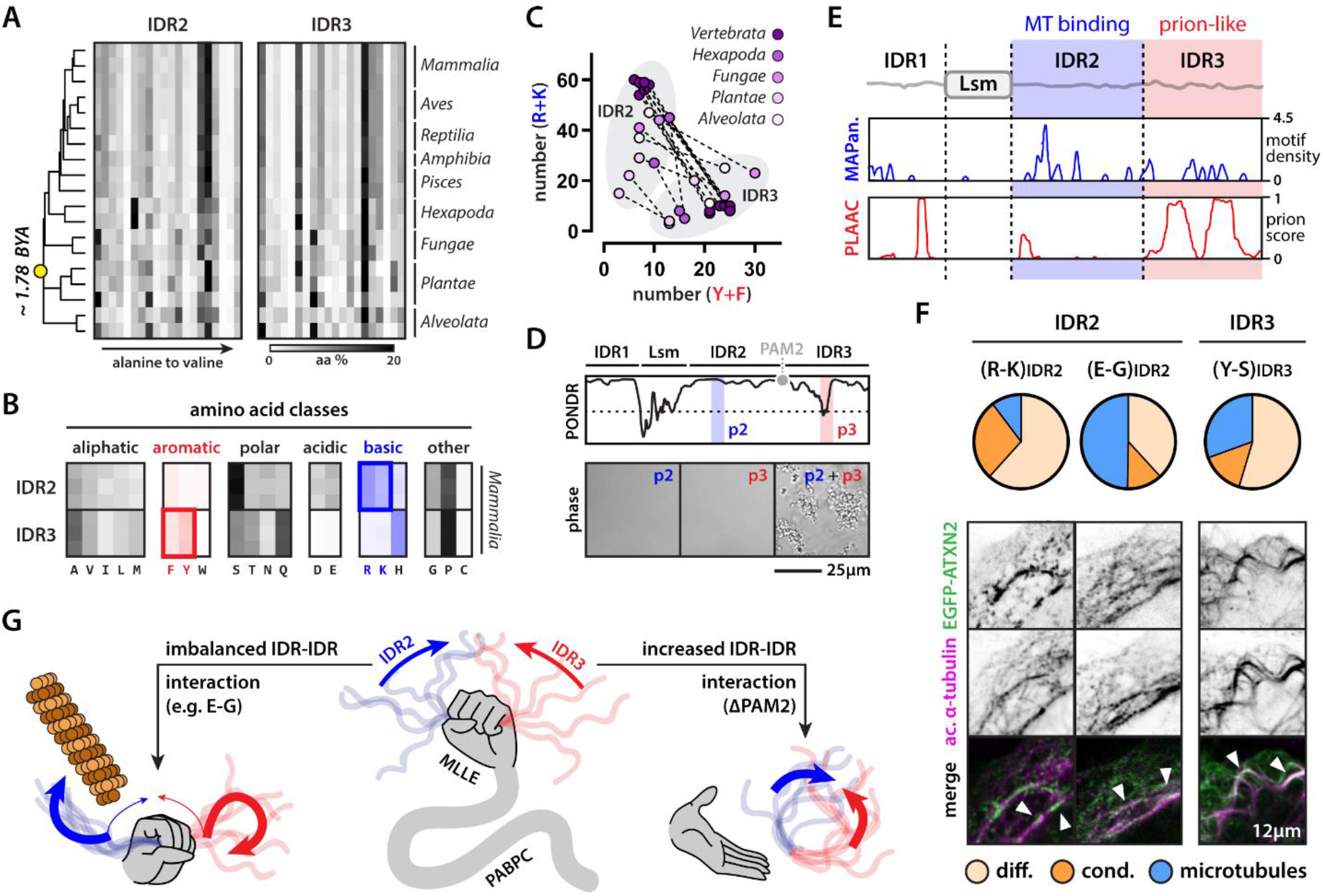
IDR2-IDR3 interaction strength regulates ATXN2 behavior. (A) Evolutionary comparison of IDR2 and IDR3 amino acid composition across the eukaryote lineage. Yellow dot highlights last common eukaryote ancestor. (B) Specific classes of amino acids are differentially enriched/depleted in IDR2 and IDR3. Only mammalian species are shown. Aromatic and basic residues are highlighted red and blue shades respectively. (C) For all tested eukaryote species, the differential IDR enrichment of basic versus aromatic residues is conserved. Dashed line connects IDR2 and IDR3 of the same species. (D) Peptides derived from IDR2 (p2) and IDR3 (p3) undergo condensation when combined. Shaded areas highlight location of peptide along the ATXN2 sequence. (E) MAPanalyzer predicts IDR2 to be a microtubule binding domain. PLAAC predicts IDR3 and the polyQ repeat to be prion-like domains. (F) Altering the pi-interaction capacity (R-K substitution) or the (relative) positive charge (E-G substitution) of IDR2, or the aromatic content (Y-S substitution) of IDR3, drives aberrant ATXN2 behavior. n = [107-157] cells (see also Fig. S1). Arrowheads highlight interaction of ATXN2 with microtubules. (G) Scheme highlighting the necessity of a balance in IDR2-IDR3 interaction strength to regulate ATXN2 behavior. Increased interaction (i.e., ΔPAM2) drives spontaneous condensation, whereas decreased interaction (e.g., E-G, ΔIDR3, etc.) promotes alternate affinities of IDR2 and drives microtubule binding. MLLE is the PAM2-binding domain of PABPC.

Having established a functional role of the PAM2 motif as a modulator of IDR2-IDR3 interaction, we next explored behavioral differences between these two domains. We performed a bioinformatics analysis, PLAAC (Alberti et al., 2009), which predicts that IDR3 is a prion-like domain (Fig. 3E). Prion-like domains can drive the condensation of a wide array of proteins across the tree of life (Newby and Lindquist, 2013). This is consistent with our earlier findings that ΔIDR3 mutants are unable to spontaneously condense, even when missing the PAM2 motif (Fig. 1J). But it remained unclear why these mutants displayed overt microtubule binding (Fig. 1J). Surprisingly, when using MAPAnalyzer –a prediction software trained on experimentally verified microtubule-binding proteins (Zhou et al., 2015)– ATXN2 is predicted to be a high confidence hit. More specifically, this prediction seems to be driven by an accumulation of short linear motifs commonly found in such proteins in the ATXN2 IDR2 domain (Fig. 3E). Indeed, deleting IDR2 completely prevented microtubule localization of ATXN2 (Fig. 1J). Fitting this data into our model (Fig. 3G), we hypothesized that weakening the interaction between IDR2 and IDR3 drives alternate affinities of these IDRs, such as microtubule binding for IDR2. To test this hypothesis, we generated amino acid substitutions to IDR2 and IDR3 to tune their affinity for one another. We first lowered the affinity of IDR2 for IDR3 by substituting its arginines for lysines. Both residues are positively charged but only arginines can effectively engage in pi-pi interactions with aromatic groups (Gallivan and Dougherty, 1999). This increased the proportion of cells with spontaneous condensates, which colocalized with the microtubule cytoskeleton (Fig. 3F, Fig. S1B). Second, we increased the relative positive charge of IDR2 by substituting glutamic acid residues for glycines. This perturbation creates an excess in positive charge, which is no longer balanced by the aromatic residues in IDR3. Indeed, the (E-G)IDR2 mutant showed increased microtubule binding (Fig. 3F, Fig. S1B). Third, substituting tyrosines for serines in IDR3 strongly reduced the affinity of this domain for IDR2, which again resulted in increased microtubule binding for this mutant (Fig. 3F, Fig. S1B).

We initially set out to understand the rules governing ATXN2’s condensation behavior. By systematically dissecting the individual domain contributions and their interactions, we uncovered a hierarchy of *cis* (IDR2-IDR3) and *trans* (PAM2-PABPC and RNA-LSM) interactions, whose balance tunes ATXN2’s solubility (Fig. 3G).

### PABPC is a holdase

Our findings indicate that ATXN2 requires the binding of PABPC to its C-terminal IDR, comprised of IDR2 and IDR3, to remain soluble. This suggests the possibility that PABPC functions as a holdase. Holdases are chaperones that bind to unfolded or disordered proteins to prevent their condensation (Hall, 2020). To directly test the hypothesis that PABPC functions as a holdase, we performed experiments using orthogonal designer condensates. We recently developed the PopTag system, a modular platform for generating synthetic and orthogonal condensates in human cells (Lasker et al., 2021). This condensation module can be functionalized with a variety of fusion proteins. When fused to GFP, we create fluorescent condensates in the cytoplasm. These condensates can then be fused to “probes” to drive specific protein-protein interactions (Fig. 4A). The ATXN2-PABPC interaction is mediated by the PAM2 motif and the MLLE domain (Jimenez-Lopez and Guzman, 2014; Kozlov et al., 2010; Xie et al., 2014). Whereas standard PopTag (GFP-Pop) condensates exclude ATXN2, fusing them to the MLLE probe drives the recruitment of ATXN2 into MLLE-PopTag condensates (Fig. 4B). Likewise, standard PopTag condensates exclude PABPC but fusing them to the PAM2 motif results in the recruitment of PABPC to these condensates (Fig. 4C). Importantly, the PAM2-PopTag was no longer able to coalesce into large condensates, but instead formed numerous small granules (Fig. 4C), reminiscent of what we had seen for wildtype and ΔPAM2 *Co*ATXN2 in *C. owczarzaki* (Fig. 2B). We confirmed that MLLE-PopTag, but not PAM2-PopTag, condensates recruit PAM2-containing proteins (Fig. 4D-E), including ATXN2 and NFX1, and that only PAM2-PopTag but not MLLE-PopTag, condensates recruit PABPC (Fig. 4F), highlighting the specificity of our system. Together, two observations support the dual role of the ATXN2-PABPC interaction (Fig. 4G) and provide evidence that PABPC can function as a *bona fide* holdase for ATXN2. First, ATXN2 is recruited into MLLE-containing condensates via its PAM2 motif, and these can be physiological PABPC-positive SGs (Fig. 1E) or synthetic MLLE-PopTag condensates (Fig. 4B). Second, PABPC binding counteracts condensation of PAM2-containing proteins, like ATXN2 (Fig. 1G) and PAM2-PopTag (Fig. 4C).

**Figure 4:**
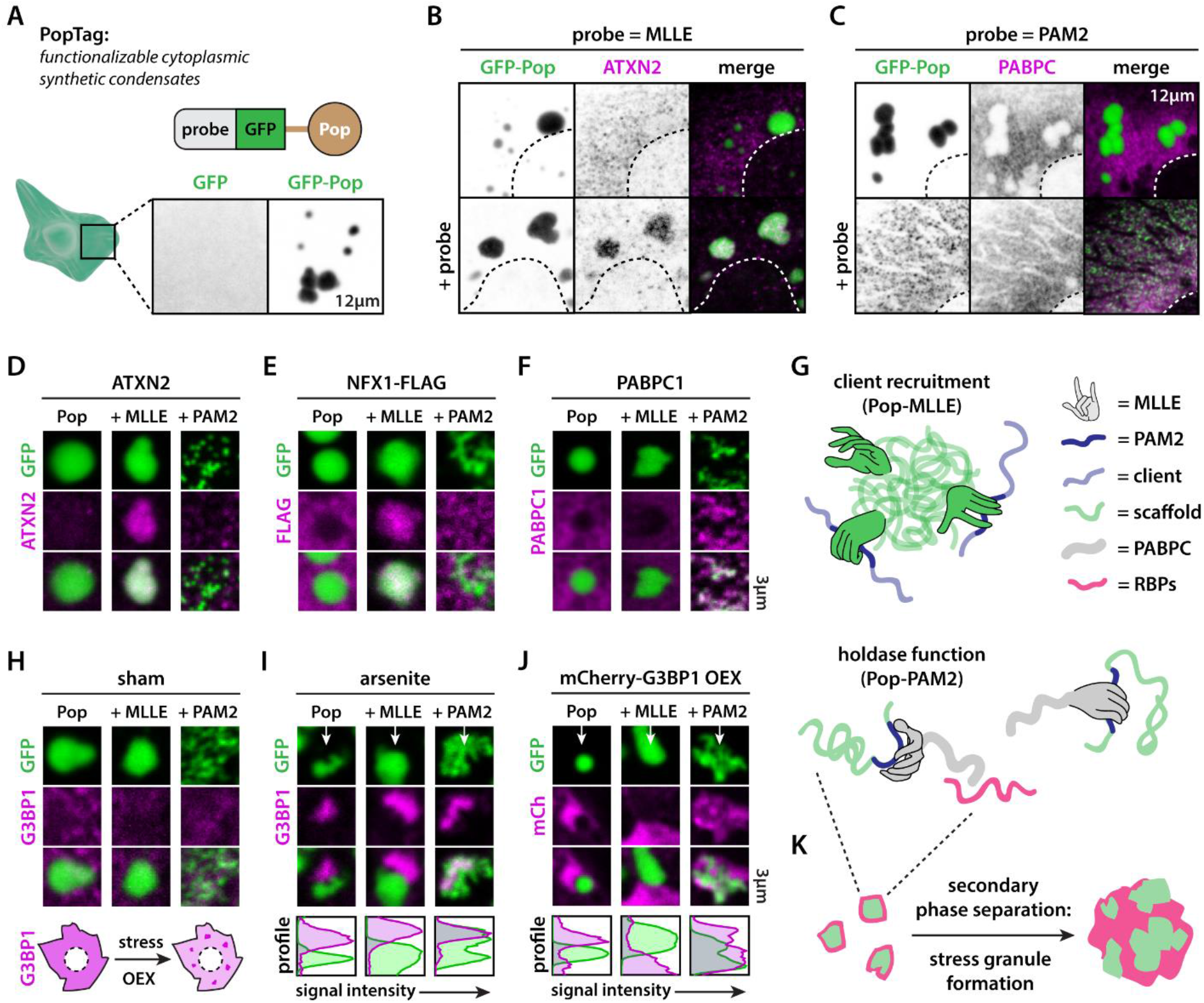
PABPC acts as a holdase and emulsifies immiscible condensates. (A) Scheme illustrating the design and use of the PopTag system. Fusing the PopTag to proteins drives their cytoplasmic condensation. (B) Functionalizing PopTag condensates with the PABPC1-derived MLLE domain drives ATXN2 partitioning into PopTag condensates. (C) Fusing GFP-PopTag to the ATXN2-derived PAM2 motif recruits PABPC1 and prevents formation of large PopTag condensates. (D) Only MLLE-PopTag fusions recruit ATXN2. (E) Only MLLE-PopTag fusions recruit the PAM2-containing protein NFX1. (F) Only PAM2-PopTag fusions recruit PABPC1 and prevent coalescence of small PopTag granules into larger condensates. (G) Scheme highlighting the effect of functionalizing synthetic PopTag condensates with the PAM2 motif or MLLE domain. (H) G3BP1 is not recruited to PopTag condensates under non-stress conditions. (I-J) Arsenite stress or G3BP1 overexpression drive stress granule formation. Small preformed PAM2-PopTag granules coalesce into larger condensates that mix with G3BP1-positive stress granules Other PopTag condensates do not mix with stress granules. (K) Scheme highlighting the incorporation of preformed PAM2-PopTag condensates into stress granules via PABPC1-mediated protein or RNA interactions. RNA not shown in scheme for clarity.

### PABPC is a condensate emulsifier

We next used our synthetic system to investigate the mechanism by which PABPC regulates protein phase separation. PABPC is a key scaffold component of cytoplasmic mRNA granules where it oligomerizes on poly(A)-tails and recruits an array of effector proteins involved in translational regulation (Goss and Kleiman, 2013). Under times of stress, ribosomes will dissociate from these mRNA granules, which causes them to condense into SGs. An emerging view is that there are preformed “cores” that condense into mature SGs driven via liquid-liquid phase separation of other proteins (Jain et al., 2016; Wheeler et al., 2016). One example of these kinds of scaffold proteins is the essential SG protein G3BP1 and its paralog G3BP2 (Kedersha et al., 2016; Yang et al., 2020). Under standard conditions, G3BP1 did not interact with any of our designer condensates (Fig. 4H). When we induced G3BP1 phase separation, via cellular stress or overexpression (Fig. 4I-J), we observed that while G3BP1 SGs do not interact with PopTag and MLLE-PopTag condensates, they do condense together with the PAM2-PopTag granules. This suggests that PABPC can drive the condensation of preformed granules aided by the phase separation of an additional RNA-binding protein, and is a synthetic postulate of the core-shell hypothesis that has been formulated to explain SG biogenesis (Fig. 4K) (Jain et al., 2016; Wheeler et al., 2016). Furthermore, since PopTag condensates themselves do not have an affinity for SGs, PABPC effectively acts as an emulsifier of immiscible condensates. As such, we propose that the mixing of ATXN2 into SGs can be considered the result of the PAM2-mediated interaction of immiscible condensates (i.e., ΔPAM2 ATXN2 granules) with PABPC acting as the emulsifier (Fig. 1E-F).

### RNA-binding is required for PABPC to function as a holdase and emulsifier

Using natural and synthetic condensate systems, we have provided evidence that PABPC has holdase and emulsifier activities. We next sought to define the mechanisms by which it exerts these functions. Because there are multiple compensatory paralogs (Xie et al., 2021) and PABPC’s function in translational control is essential for normal cell physiology (Gorgoni et al., 2011), we generated a designer holdase system to circumvent these potentially confounding factors. As we have shown above, PABPC recognizes its clients via the PAM2-MLLE interaction (Fig. 5A). By simply substituting this natural interaction pair for a synthetic version (Fig. 5B), in this case the HA-tag and an engineered HA-binding nanobody called Frankenbody (F-body) (Zhao et al., 2019), we can disentangle the effects of endogenous PABPC from engineered ones. Whereas PAM2-PopTag cannot coalesce into large condensates, HA-PopTag can (Fig. 5C). This property allows us to introduce designer condensates and evaluate their activity (Fig. S3A-B). mCherry was not recruited into HA-PopTag condensates but mCherry-tagged F-body was strongly recruited to HA-PopTag condensates (Fig. 5D). However, this protein was not able to prevent the formation of large condensates. On the other hand, F-PABPC, which is full-length PABPC1 with its MLLE domain replaced by the F-body (Fig. 5E), completely prevented HA-PopTag coalescence (Fig. 5D, Fig. S3C). Since we were able to reconstitute PABPC holdase activity in a synthetic system, we could now start interrogating the role of underlying molecular features. We engineered an RNA-binding deficient mutant by substituting key aromatic residues in the RRMs for leucines (Buratti and Baralle, 2001), called F-PABPC*. This mutant was unable to efficiently prevent HA-PopTag condensation, indicating that RNA binding is important for PABPC’s holdase activity (Fig. 5D, Fig. S3C). To test whether poly(A) binding was specifically required, or whether RNA binding in general could confer this activity, we engineered holdase versions where we substituted the four PABPC RRM domains for other tetravalent RNA binding domains (hnRNPI, MBNL1 and FXR1) (Fig. 5E). Synthetic holdases carrying the hnRNPI or MBNL1 RNA-binding domains were active but the FXR1 chimera was not (Fig. 5F-G, Fig. S3C). Thus, PABPC holdase activity depends on specific RNA-protein interactions or RNA species.

**Figure 5:**
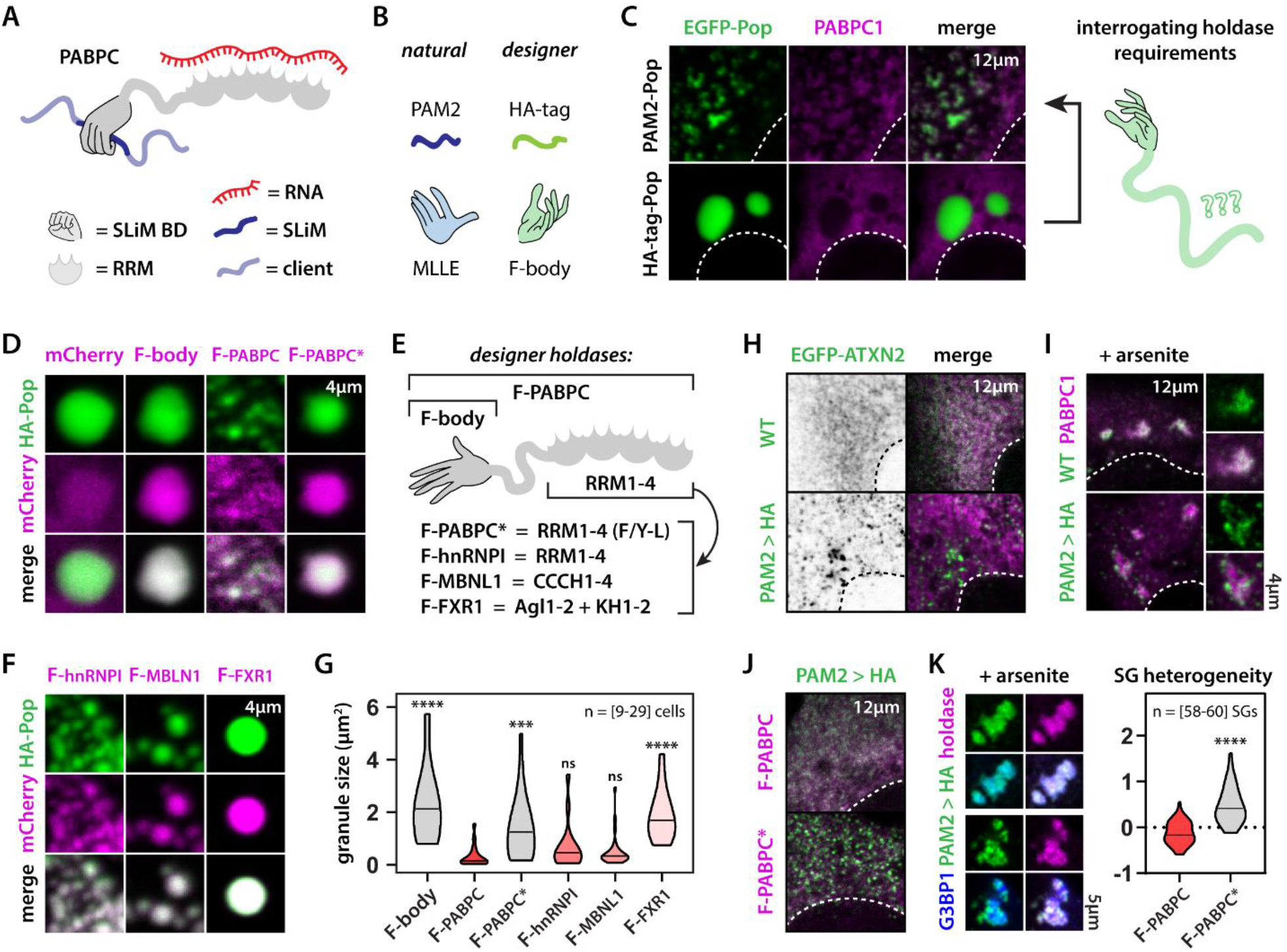
RNA binding is required for PABPC holdase activity. (A) Scheme highlighting PABPC domain structure. (B) Examples of a natural and synthetic SLiM-based interaction pair. (C) PAM2-PopTag fails to condense into large condensates due to PABPC interaction. HA-PopTag condensates do not interact with endogenous PABPC. This allows us to interrogate sequence requirements of holdase activity using designer holdases. (D) mCherry is diffusely localized throughout the cytoplasm and Ha-PopTag condensates. F-body strongly partitions into HA-PopTag condensates. F-PABPC binds to HA-PopTag granules and prevents their coalescence into large condensates, whereas this was not the case for the RNA-binding deficient F-PABPC* mutant. (E) Scheme highlighting domain architecture of designer condensates and RNA-binding mutants. (F) Holdase activity is dependent on the involved RNA-binding domains. (G) Quantification of activity of designer holdases. Cells combined from three experiments (see also Fig. S3). One-way ANOVA. (H) Replacing the PAM2 motif with an HA-tag drives the spontaneous condensation of ATXN2. (I) PAM2>HA ATXN2 fails to properly partition into PAPC stress granules. (J-K) F-PABPC but not F-PABPC* rescues spontaneous condensation (J) and stress granule demixing (K) of PAM2>HA ATXN2. Mann-Whitney. *** p-value < 0.001, **** p-value < 0.0001.

The results of our PopTag system provide evidence that RNA binding is required for PABPC’s holdase activity. To test if these findings also applied to ATXN2, we first created a mutant where we replaced the PAM2 motif with the HA-tag. Similar to what we observed for ΔPAM2, this mutant spontaneously condensed under standard conditions and was unable to properly partition into SGs (Fig. 5H-I). Co-expressing our engineered holdases revealed that RNA-binding by F-PABPC is also required to prevent spontaneous condensation and to drive SG partitioning of the HA-tag mutant ATXN2 (Fig. 5J-K). Thus, our data indicate that PABPC is a holdase regulating ATXN2 protein phase separation by virtue of its interaction with RNA.

## DISCUSSION

SGs have been extensively studied as model biomolecular condensates (Alberti et al., 2017; Boeynaems et al., 2018; Protter and Parker, 2016; Shin and Brangwynne, 2017; Wolozin and Ivanov, 2019; Zhang et al., 2019). But an understanding of the complex molecular interactions that govern their behavior in health and disease is still incomplete. How hundreds of different proteins specifically partition into these assemblies, and the functional implications of this, remain largely unknown. Since several SG proteins have both RNA-binding and sticky disordered domains, an initial hypothesis was that this combination was sufficient to drive partitioning of a protein into these condensates. Here we focused on ATXN2 as a case study of a disordered RNA-binding protein, and surprisingly found that its SG targeting is exclusively mediated by a small 14aa SLiM that engages PABPC. Such motifs are abundant in the proteome (Tompa et al., 2014) and have been previously implicated in the regulation of SG proteins, as sites of post-translational modifications (Monahan et al., 2017) or as localization sequences for nucleocytoplasmic transport (Dormann et al., 2012). However, this is to our knowledge the first motif that can specifically drive SG partitioning. Similar to karyopherins, which mediate SLiM-based nuclear import but have been recently shown to counteract (aberrant) protein condensation (Guo et al., 2018; Hofweber et al., 2018; Qamar et al., 2018), PABPC also seems to moonlight as an unconventional holdase. Holdases are chaperones that maintain disordered or misfolded clients in a specific conformation, preventing their condensation (Hall, 2020). We found that PABPC modulates IDR2-IDR3 interactions that otherwise drive ATXN2 condensation. Via a synthetic biology approach making use of designer condensates and engineered holdases, we have provided evidence that PABPC acts as a general holdase regulating the condensation of natural or synthetic PAM2 proteins, and that this activity is dependent on RNA binding. Moreover, we find that this chaperone-client interaction between PABPC and ATXN2 is functionally conserved across eukaryotes, highlighting the biological importance of this ancient regulatory switch.

PABPC prevents the overt interaction of ATXN2 IDRs but we find that their interaction is important for regulating normal ATXN2 behavior. Just as for other SG proteins, ATXN2 contains IDRs that are differentially enriched in basic or aromatic residues, which can engage one another via pi-pi and cation-pi interactions (Wang et al., 2018). Since it is exactly the interplay between such domains that seems to drive the neurotoxicity of these proteins, one question remains as to why this combination of extremely sticky domains is so prevalent? Unexpectedly, when we weakened the affinity of the ATXN2 IDRs for one another, we found that this resulted in aberrant behavior of the protein. Specifically, we observed the spontaneous coating of the microtubule cytoskeleton by these ATXN2 mutants. Indeed, bioinformatics analysis predicted the basic IDR2 to act as a microtubule-binding domain. Therefore, it seems that the ATXN2 prion-like domain (IDR3) effectively quenches the microtubule affinity of IDR2, facilitating its normal behavior. Intriguingly, ATXN2 has been implicated in the regulation of the microtubule cytoskeleton in both *Drosophila* (del Castillo et al., 2021) and *Caenorhabditis* (Gnazzo et al., 2016; Stubenvoll et al., 2016), with one study showing its enrichment on the mitotic spindle (Gnazzo et al., 2016). While it remains unknown if ATXN2 functions similarly in mammalian systems, it is noteworthy that ATXN2 and other SG proteins have been found in several mass spectrometry datasets of microtubule-binding proteins (Fig. S4). Moreover, since several of the implicated proteins carry both basic and aromatic IDRs, it will be interesting to see if such prion-like domains can more broadly act as quenchers of basic IDRs, thereby regulating their affinities for specific interaction partners. In this way, basic disordered peptides produced by abnormal translation of expanded repeats, have now been identified in three independent degenerative conditions (Mizielinska et al., 2014; Todd et al., 2020; Zu et al., 2017). These peptides show extreme toxicity in an array of disease models and can broadly dysregulate cellular metabolism (Odeh and Shorter, 2020). Notably, such basic peptides potently interact with both prion-like proteins (Boeynaems et al., 2017a) and the microtubule cytoskeleton (Fumagalli et al., 2021). Additionally, translation readthrough events have been recently found to result in the generation of poly-K repeat extensions tails by translation of the mRNA’s poly(A)-tail. Such poly-K modified proteins were found to target and perturb nucleoli (Davis et al., 2021). These “biological accidents” that lead to the uncontrolled expression of an unquenched basic IDR (Kwon et al., 2014), show how important the correct regulation of these sticky sequences is to normal cell physiology.

Our data indicates that the precise balance between *cis* (i.e., IDR2-IDR3) and *trans* (i.e., RNA and PABPC) interactions tunes ATXN2’s behavior. However, the dependence of this system on the client-holdase interaction may form its Achilles’ heel in disease: polyQ expansion drives irreversible ATXN2 aggregation in patients and disease models (Alves-Cruzeiro et al., 2016). Despite chaperoning the C-terminal IDR against spontaneous condensation, PABPC is not able to counteract aggregation caused by the N-terminal polyQ domain. Pathology data from murine models has shown that PABPC colocalizes with the ATXN2 inclusion bodies (Damrath et al., 2012; Sen et al., 2019). Since PABPC serves an important role in translational regulation, its sequestration into these irreversible aggregates could affect neuronal health via its loss of function. Such a mechanism would explain previous results from a *Drosophila* ATXN2 model that required the PAM2-mediated interaction with PABPC for neurotoxicity (Kim et al., 2014). This finding illuminates a new aspect of SCA2 and ALS pathogenesis, and could inspire novel opportunities for therapeutic intervention (e.g., designer chaperones).

In conclusion, by using a multidisciplinary approach involving evolutionary analyses combined with synthetic biology, we uncover a conserved and unexpected novel function for a major RNA-binding protein. Additionally, our results highlight how a complex hierarchy of *cis* and *trans* interactions mediate the precise behavior of stress granule proteins.

## ACKNOWLEDGEMENTS

We thank all members of the Gitler lab as well as the Carnegie-Stanford Intrinsically Disordered Protein Scientific Interest Group (IDPSIG) for helpful discussion and suggestions. We thank the Stanford Neuroscience Microscopy Service for use of the core facility. We thank Dr. Seung Y. Rhee for resources and feedback, G. Materassi-Shultz for plant growth facilities management, and the Carnegie Advanced Imaging facility.

## AUTHOR CONTRIBUTIONS

**S.B:** Conceptualization, Methodology, Validation, Formal Analysis, Investigation, Data Curation, Writing-Original Draft, Visualization. **Y.D**.: Investigation. **A.M**.: Investigation, Formal Analysis. **V.S**.: Investigation. **G.H**.: Investigation. **G.K**.: Resources. **A.S**.: Investigation. **N-E.S**.: Investigation. **R.D**.: Supervision. **I.R-T**.: Supervision. **K.L**.: Methodology. **G.A**.: Supervision. **E.K**.: Supervision. **A.D.G**.: Conceptualization, Supervision, Funding acquisition, Writing-Review & Editing.

## SUPPLEMENTAL INFORMATION

## OTHER SUPPLEMENTAL MATERIAL

Table S1 (Excel): Sequences of protein mutants, orthologs and peptides used in this study.

## STAR METHODS

### RESOURCE AVAILABILITY

#### Lead contact

Further information and requests for resources and reagents should be directed to and will be fulfilled by the lead contact, Aaron D. Gitler.

#### Materials availability

All unique materials generated in this study will be made available on request from the Lead Contact upon completion of a Materials Transfer Agreement.

#### Data and code availability

The published article includes all datasets generated or analyzed during this study.

### EXPERIMENTAL MODEL AND SUBJECT DETAILS

#### Human cell lines

U2OS (ATCC, HTB-96) and HeLa cells (ATCC, CCL-2) were grown at 37°C in a humidified atmosphere with 5% CO2 for 24h in Dulbecco’s Modified Eagle’s Medium (DMEM), high glucose, GlutaMAX + 10% Fetal Bovine Serum (FBS) and pen/strep (Thermo Fisher Scientific).

##### Capsapora owczarzaki

*C. owczarzaki* was grown axenically in cell culture at 23°C in ATCC medium 1034 (modified PYNFH medium).

##### Trypanosoma brucei

*T. brucei* PCF 29–13 (T7RNAP NEO TETR HYG) co-expressing T7 RNA polymerase and Tet repressor was grown in SDM-79 medium as previously described (Huang et al., 2013), supplemented with hemin (7.5 μg/ml) and 10% heat-inactivated fetal bovine serum and at 27°C in the presence of G418 (15 μg/ml) and hygromycin (50 μg/ml) to maintain the integrated genes for T7 RNA polymerase and tetracycline repressor, respectively.

#### Plant growth conditions

*Nicotiana benthamiana* (tobacco) plants and *Arabidopsis thaliana* plants used for seed propagation were grown in soil (PRO-MIX® HP Mycorrhizae) inside growth chambers held at 22 °C with a 16/8 hour photoperiod (130 μmol.m^-2^.s^-1^). *Arabidopsis thaliana* seedlings used for microscopy were grown on Murashige and Skoog (MS) medium (0.5X Murashige and Skoog basal salt mixture (PhytoTechnologies Laboratories) at a pH of 5.7 supplemented with 0.8 % agar (Difco) and 1 % sucrose (Sigma-Aldrich)) inside growth cabinets (Percival) held at 22 °C with a 16/8 hour photoperiod (130 μmol.m^-2^.s^-1^). Their seeds were first sterilized in 70 % ethanol by vortexing for 5 minutes followed by replacement of the solution with 100 % ethanol. Seeds were then immediately placed on pre-sterilized filter papers (Grade 410, VWR) and left to dry in a laminar flow hood. They were then sown on square petri dishes (120 × 120 wide × 15 mm high (VWR)) containing 40 mL of MS medium. Plates were then sealed with micropore surgical tape (3M) and covered in aluminum foil before being placed at 4°C for three days to break seed dormancy.

### METHOD DETAILS

#### Plasmid construction

Sequences of all mutants, orthologs, and holdases are shown in Table S1.

Constructs for human expression were generated through custom synthesis and subcloned into a pcDNA3.1 backbone by Genscript (Piscataway, USA). The G3BP1-mCherry construct was a kind gift of Dr. Nancy Kedersha and Dr. Paul Anderson (Harvard Medical School, USA).

Constructs for *Co*ATXN2 (CAOG_07908) were synthesized by Genscript and subcloned into the pONSY vector (Addgene, 111873) using Gibson assembly (Gibson et al., 2009). Prior to assembly, the backbone vector and the insert were prepared as following: the pONSY vector was linearized with EcoRV-HF restriction enzyme (New England BioLabs); the cDNA coding for ATXN2-GFP or dPAM-ATXN2-GFP was amplified using Phusion High-Fidelity DNA Polymerase (New England BioLabs) with the forward and reverse primers (see below). TOP10 E. coli cells were transformed with the product of the assembly reaction and positive clones were confirmed by Sanger sequencing.

Forward primer *Co*ATXN2: CGGGACTAGTGATATCATGAGCAAGGGCGAGGAG

Reverse primer *Co*ATXN2: GCAAACACAAAATTCAAACGGGCCCTGCCTT

Constructs for *Tb*ATXN2 (Tb927.8.4540) were synthesized by Genscript and subcloned into the *T. brucei* expression vector pLew100v5_bsd to generate pLew100 (*GFP-TbATXN2*) and pLew100(*GFP-dPAM-TbATXN2*) (Huang et al., 2013). Prior to subcloning, the backbone vector and the insert were prepared as following: the pLew100v5_bsd vector was digested with *Xba*I and *Bam*HI restriction enzymes (New England BioLabs) and gel-purified; the cDNA coding for GFP-*Tb*ATXN2 or GFP-dPAM-*Tb*ATXN2 was amplified using Q5 High-Fidelity DNA Polymerase (New England BioLabs) with forward and reverse primers (see below, digested with *Xba*I and *Bam*HI, and purified. The enzyme-cut pLew100v5_bsd and GFP-*Tb*ATXN2 or GFP-dPAM-*Tb*ATXN2 were ligated and transformed into DH5α *E. coli* cells, and the positive clones were confirmed by Sanger sequencing at Genewiz (Research Triangle Park, NC).

Forward primer *Tb*ATXN2: 5’GC<u>TCTAGA</u>TAAGGCACCATGAGCAAGGGCGAGGAGCTG-3’ (*<u>Xba</u>*<u>I site</u>)

Reverse primer *Tb*ATXN2: 5’CG<u>GGATCC</u>CTATTTCCCAACTCGTTTCTTCGGCC-3’ (*<u>Bam</u>*<u>HI site</u>)

Constructs for *At*ATXN2 (AT3G14010) and ΔPAM2 *At*ATXN2 were synthesized by GenScript Biotech Corporation (Piscataway, NJ) with flanking Gateway attB sites. They were then BP recombined using the Gateway system (Thermo Fisher Scientific) into pDONR221, and then subcloned using LR recombination (Thermo Fisher Scientific) into pGWB606 (https://shimane-u.org/nakagawa/pgwb-tables/4.htm) (Nakagawa et al., 2007) to generate p35S:GFP-*At*ATXN2 and p35S:GFP-ΔPAM2AtATXN2. To generate p35S:RFP-PAB2, a pENTR223 vector (GC104970) containing PAB2’s cDNA (AT4G34110) was first obtained by the *Arabidopsis* Biological Resource Center (ABRC). Since it contained the same resistance, spectinomycin, as the destination vector pGWB661 (https://shimane-u.org/nakagawa/pgwb-tables/4.htm) (Nakagawa et al., 2007), it was first subcloned into pDEST15 (Thermo Fisher Scientific) using LR recombination (Thermo Fisher Scientific) and then subcloned into another entry vector, pDONR221, (Thermo Fisher Scientific) using BP recombination (Thermo Fisher Scientific). Finally, the PAB2 cDNA sequence was then subcloned into pGWB661 to generate p35S:RFP-PAB2.

#### Human cell culture, treatments and microscopy

U2OS and HeLa cells were grown at 37 °C in a humidified atmosphere with 5 % CO2 for 24 h in Dulbecco’s Modified Eagle’s Medium (DMEM), high glucose, GlutaMAX + 10 % Fetal Bovine Serum (FBS) and pen/strep (Thermo Fisher Scientific). Cells were transiently transfected using Lipofectamine 3000 (Thermo Fisher Scientific) according to manufacturer’s instructions. Cells grown on cover slips were fixed for 24 h after transfection in 4 % formaldehyde in PBS. Slides were mounted using ProLong Gold antifade reagent (Life Technologies). Confocal images were obtained using a Zeiss LSM 710 confocal microscope. Images were processed and analyzed using Fiji (Schindelin et al., 2012). To induce stress granule formation, cells were treated for 1 hour with 250 µM of sodium arsenite (Sigma-Aldrich).

#### FRAP measurements in human cells

U2OS cells were cultured in glass bottom dishes (Ibidi) and transfected with GFP-ATXN2 constructs as described above. After 24 h, GFP-ATXN2 condensates were bleached and fluorescence recovery after bleaching was monitored using ZEN software on a Zeiss LSM 710 confocal microscope with incubation chamber at 37 °C and 5 % CO2. Data were analyzed as described previously (Boeynaems et al., 2017b). In brief, raw data were background subtracted and normalized using Excel, and subsequently plotted using GraphPad Prism 8.4.1 software.

#### ATXN2 KO generation in human cells

An ATXN2-targeting sgRNA sequence (GATGGCATGGAGCCCCGATCC) was cloned into a lentiviral backbone containing mCherry and puromycin resistance cassette. We then transfected this construct into HEK293T cells (from ATCC) at 70-80% confluency in 6-well plates. The resulting supernatant, collected 48 hours later using a syringe through a 0.45um filter (EMD Millipore; SLHP033RS), was used to infect low-passage HeLa-cas9 cells (HeLa cells expressing cas9) for generation of the knockout line. HeLa-Cas9 cells were cultured in DMEM containing high glucose, 10% FBS, and pen/strep in a 10cm plate. Virus titering was performed such that MOI of the sgRNA sequence-containing construct was <40% (as determined by % of cells that were mCherry-positive). The media was changed 24 hours after infection to get rid of lentivirus-containing media. One week after infection, mCherry-positive cells (cells that incorporated a sgRNA) were single-cell sorted into a 96-well plate. Clones were grown up then Sanger sequenced at the ATXN2 locus to determine successful knockout. One of the confirmed clonal knockout lines was used for this study.

#### *Capsaspora* cell transfection and microscopy

Confluent *Capsaspora* cells were co-transfected with the pONSY-H2B-mCherry and pONSY-ATXN2-GFP or pONSY-dPAM2-ATXN2-GFP plasmids as described elsewhere (Parra-Acero et al., 2018). For life imaging, cells were seeded in a µ-Slide 4-well glass-bottom dish (Ibidi). Pictures were taken 2 days after transfection on a Zeiss Axio Observer Z.1 epifluorescence inverted microscope equipped with LED illumination and a Axioscan 503 mono camera. A 100x immersion oil objective was used. All pictures were taken at the same laser intensity and exposure settings.

#### *Trypanosoma* cell transfection, immunoblotting and microscopy

The plasmid pLew100 (*GFP-TbATXN2*) or pLew100(*GFP-dPAM-TbATXN2*) was *No*tI-linearized and transfected into mid-log phase *T. brucei* PCF, as described previously (Huang et al., 2013). The stable transformants were obtained in SDM-79 medium supplemented with 15% FBS plus the appropriate antibiotic (15 μg/ml G418, 50 μg/ml hygromycin, and 10 μg/ml blasticidin). Expression of GFP-TbATXN2 or GFP-dPAM-TbATXN2 was induced with 1 μg/ml fresh tetracycline and confirmed by immunoblotting and microscopy, as described below.

The blots were incubated with rabbit antibodies against GFP (1:2,500) or mouse antibodies against tubulin (1:10,000) for 1 h. After five washings with PBS-T, the blots were incubated with horseradish peroxidase-conjugated anti-rabbit or anti-mouse IgG (H+L) antibody at a dilution of 1:15,000 for 1 h. After washing five times with PBS-T, the immunoblots were visualized using Pierce ECL western blotting substrate according to the manufacturer’s instructions.

The tetracycline-induced trypanosomes were fixed with paraformaldehyde, adhered to poly-L-lysine-coated coverslips, permeabilized with Triton X-100, and blocked with BSA, as described previously (Huang et al., 2013). After blocking, trypanosomes were stained in 3% BSA/PBS with rabbit polyclonal antibody against GFP (1:250) for 1 h. After thoroughly washing with PBS containing 3% BSA, cells were incubated with Alexa 488-conjugated goat anti-rabbit antibody at 1:1,000 for 1 h. The cells were counterstained with 4′,6-diamidino-2-phenylindole (DAPI) before mounting with Gold ProLong Gold antifade reagent (Molecular Probes). Differential interference contrast (DIC) and fluorescent optical images were captured using an Olympus IX-71 inverted fluorescence microscope with a Photometrix CoolSnap^HQ^ charge-coupled device camera driven by DeltaVision software (Applied Precision, Seattle, WA). Images were deconvolved for 15 cycles using SoftwoRx deconvolution software.

#### Transgenic *Arabidopsis thaliana* lines

Transgenic plants were generated using *Agrobacterium tumefaciens*-mediated (GV3101 strain) transformation (Clough, 2005) of Col-0 with the constructs described in the *Plant plasmid construction* section. Transgenic seedlings (T_1_) were selected with Basta and left to self to generate T_2_ seeds. These were then selected on MS medium supplemented with Basta to select only those that carried one T-DNA construct as determined by the Mendelian segregation ratio (3:1) of the Basta-resistance trait. Basta-resistant seedlings from the selected lines were transferred to soil and left to self. Only lines with 100% Basta-resistant progeny (T_3_), indicating their homozygosity, were used for microscopy experiments

#### Tobacco transient assays

*Agrobacterium* lines (GV3101 strain) carrying p35S:GFP-AtATXN2, p35S:GFP-ΔPAM2AtATXN2 and p35S:RFP-PAB2, were grown overnight at 28 °C in LB broth (Fisher BioReagents) containing 25 mg/L rifampicin (Fisher BioReagents), 50 mg/mL gentamicin (GoldBio) and 50 mg/L spectinomycin (GoldBio). Cultures were washed four times with infiltration buffer (10 mM MgCl_2_ (omniPur, EMD), 10 mM MES (pH 5.6) (J. T. Baker), and 100 uM acetosyringone (Sigma-Aldrich)) and diluted to reach an OD_600_ of 0.8. An equal amount of cultures carrying p35S:RFP-PAB2 and p35S:GFP-AtATXN2 or p35S:GFP-ΔPAM2AtATXN2 were then pre-mixed. The mixtures were then infiltrated into 4^th^ or 5^th^ leaves from 6-week-old tobacco plants using Monoject 1mL Tuberculin Syringes (Covidien). For each pair of constructs, four individual tobacco plants were infiltrated. Three days after infiltration, small leaf squares (approximately 0.5 cm side length) were cut from the infiltrated regions and mounted between two cover slips in water. They were then either directly imaged (no stress condition) or placed in a 37 °C incubator for 30 min to induce SGs (heat shock) before imaging.

#### Plant microscopy

*Arabidopsis thaliana* seedlings and tobacco leaves were imaged at room temperature on a LEICA TCS SP8 laser scanning confocal microscope in resonant scanning mode using the Leica Application Suite X software. Seedlings or tobacco samples were mounted in water and then imaged using the HC PL APO CS2 63X/1.20 water objective. GFP and RFP fluorescence signals were detected by exciting with a white light laser at 488 nm and 555 nm, respectively, and by collecting emission from 500-550 nm and 565-615 nm, respectively, on a HyD SMD hybrid detector (Leica) with a lifetime gate filter of 1-6 ns to reduce background autofluorescence. Z-stacks (*Arabidopsis*) were collected with a bidirectional 64-line averaging while single-frame images (tobacco) were collected with a bidirectional 1024-line averaging. For colocalization experiments, samples were imaged sequentially between each line to ensure that the colocalization signals were not due to bleed-throughs. Images are representative of at least three biological replicates for each construct (tobacco) and of at least three independently generated transgenic lines (*Arabidopsis*).

#### IDR peptide condensation experiment

Peptide experiments were performed as described previously with slight modifications (Boeynaems et al., 2017b). In brief, IDR2 (581 – 640 aa) and IDR3 (1091 – 1150 aa) peptides were generated via chemical synthesis by Pepscan (Lelystad, the Netherlands). Peptides were dissolved in milli-Q water and stored at - 20°C. To test for condensation, peptides were diluted to a concentration of 100 µM in a 50 mM potassium phosphate buffer at pH 7.5. Samples were transferred to an imaging chamber (Grace Bio-Labs, Bend, USA) and imaged on a Zeiss LSM 710 confocal microscope. Sequences of the peptides shown in Table S1.

#### Evolutionary analysis

The cladogram depicted in Figure 3 and Figure S2 was generated using phyloT v2 (https://phylot.biobyte.de/) and iTOL v6 (https://itol.embl.de/)(Letunic and Bork, 2021). species list for cladograms for amino acid composition and PAM2 (from top to bottom): *Homo sapiens, Mus musculus, Bos taurus, Gallus gallus, Falco peregrinus, Alligator mississippiensis, Anolis carolinensis, Xenopus tropicalis, Latimeria chalumnae, Danio rerio, Apis mellifera, Drosophila melanogaster, Ustilago maydis, Aspergillus niger, Arabidopsis thaliana, Oryza sativa, Chlamydomonas reinhardtii, Naegleria gruberi, Toxoplasma gondii*. Amino acid percentages were calculated via ProtParam (https://web.expasy.org/protparam/)(Wilkins et al., 1999) and plotted in heatmaps via Morpheus (https://software.broadinstitute.org/morpheus/).

The cladogram in Figure 2 is modeled after (Burki et al., 2020).

### QUANTIFICATION AND STATISTICAL ANALYSIS

All data was analyzed using GraphPad Prism 8.4.1 and Excel. Statistical tests, p values, number of samples, replicates, and experiments are indicated in the figure legends.

## SUPPLEMENTAL FIGURES

**Figure S1:**
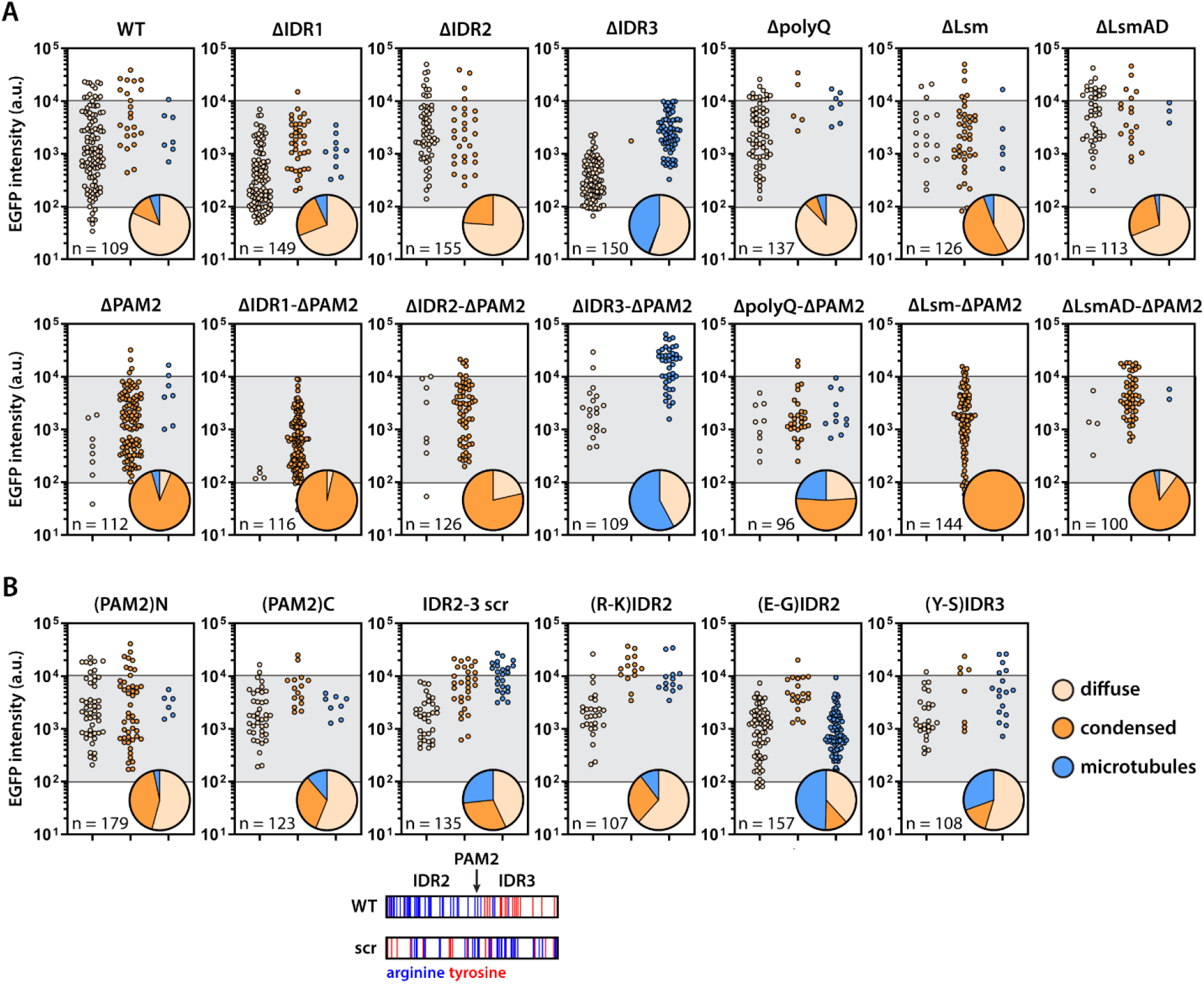
Quantification of ATXN2 behavior. (A-B) Scatterplots show cells with diffuse, condensed or microtubule-localized ATXN2. Cells combined from 3 experiments. y-axes indicate cytoplasmic EGFP intensity as a proxy for ATXN2 transgene concentration. Pie charts indicate the percentage of cells with certain behavior from the grey boxed area. This was done to correct for expression differences between ATXN2 mutants. n indicates the number of cells represented by the pie charts. (A) corresponds to mutants shown in Fig. 1J, (B) corresponds to mutants shown in Fig. 1M, Fig. 3F, and the scrambled mutant. Distribution of arginines and tyrosine is shown for the scrambled mutant.

**Figure S2:**
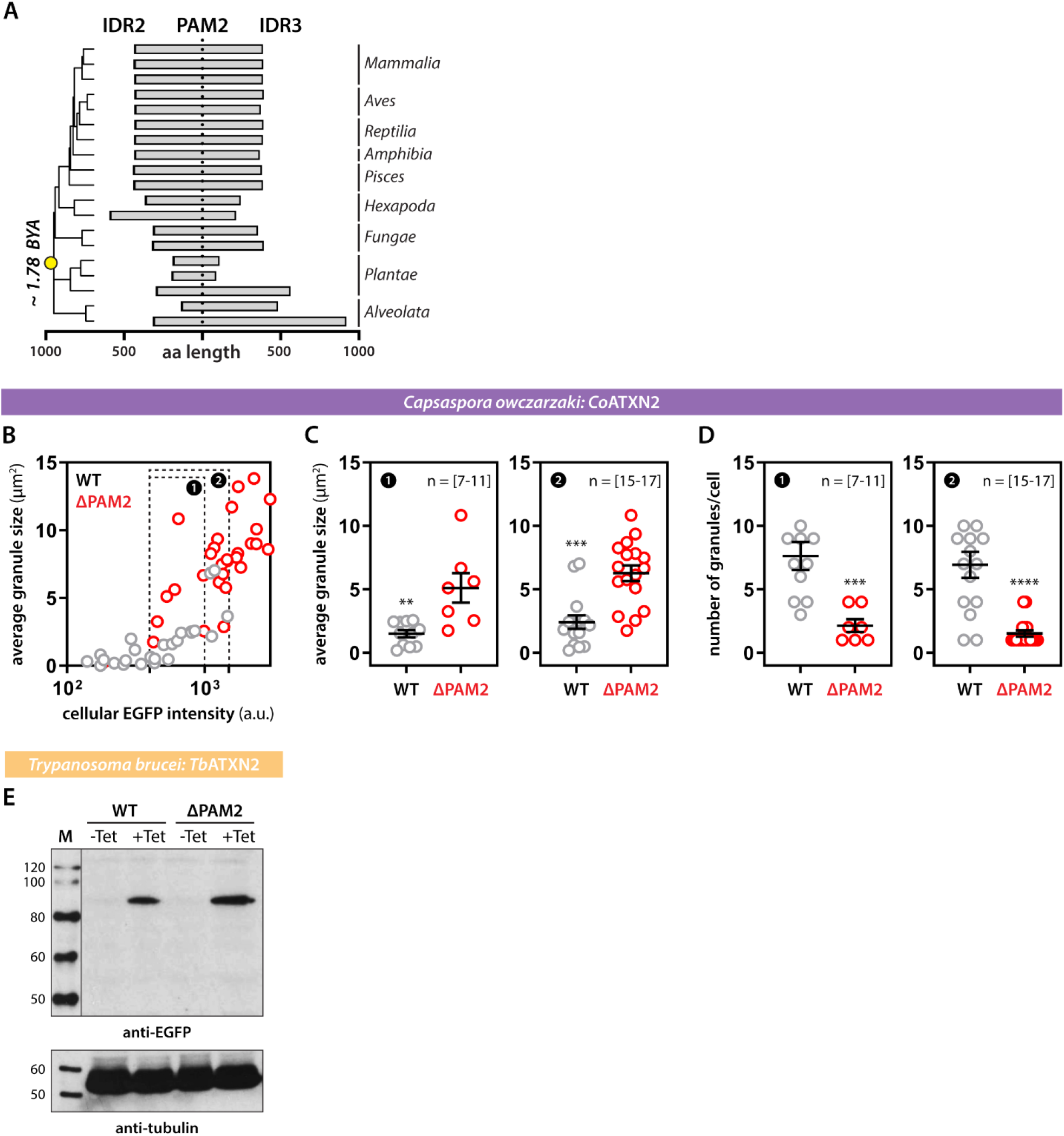
Supplementary data to Figure 2. (A) The PAM2 motif is always localized between IDR2 and IDR3 across the eukaryote lineage. Yellow dot indicates the position of the last common eukaryote ancestor. x-axis gives length information on IDR2 and IDR3, relative to the position of PAM2. Grey bars are individual proteins as in Figure 3A (B-D) Quantification of *C. owczarzaki Co*ATXN2 granules. (B) Summary of all analyzed cells. (C) Statistical analysis of average granule size per cell. (D) Statistical analysis of number of granules per cell. (1) and (2) are bins of cells with similar *Co*ATXN2 expression levels. Mann-Whitney. ** p-value < 0.01, *** p-value < 0.001, **** p-value < 0.0001. (E) Western blots show Tet-inducible expression of both WT and ΔPAM2 *Tb*ATXN2 in *T. brucei*.

**Figure S3:**
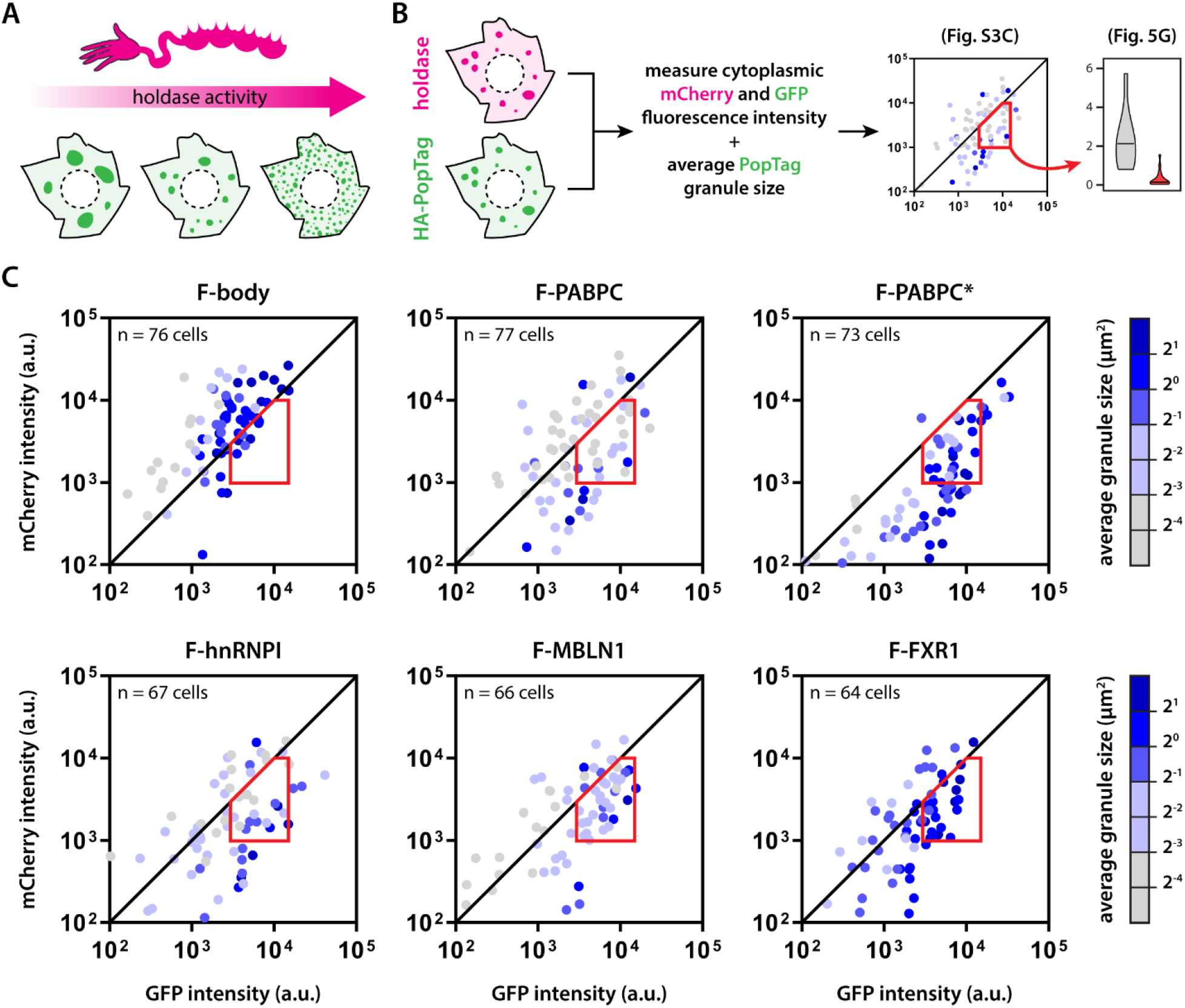
Quantification of activity of designer holdases. (A) Activity of the designer holdases correlates to PopTag granule size. (B) Scheme highlighting approach to relate granule size to expression of HA-PopTag and mCherry-holdase constructs. (C) Quantification of HA-PopTag granule size for all designer holdases. Color of each dot represents the average granule size of that cell. Cells are combined from three experiments. The red area highlights the selection of cells that were directly compared between different designer holdases (see Fig. 5G). This was done to correct for differences in expression between constructs.

**Figure S4:**
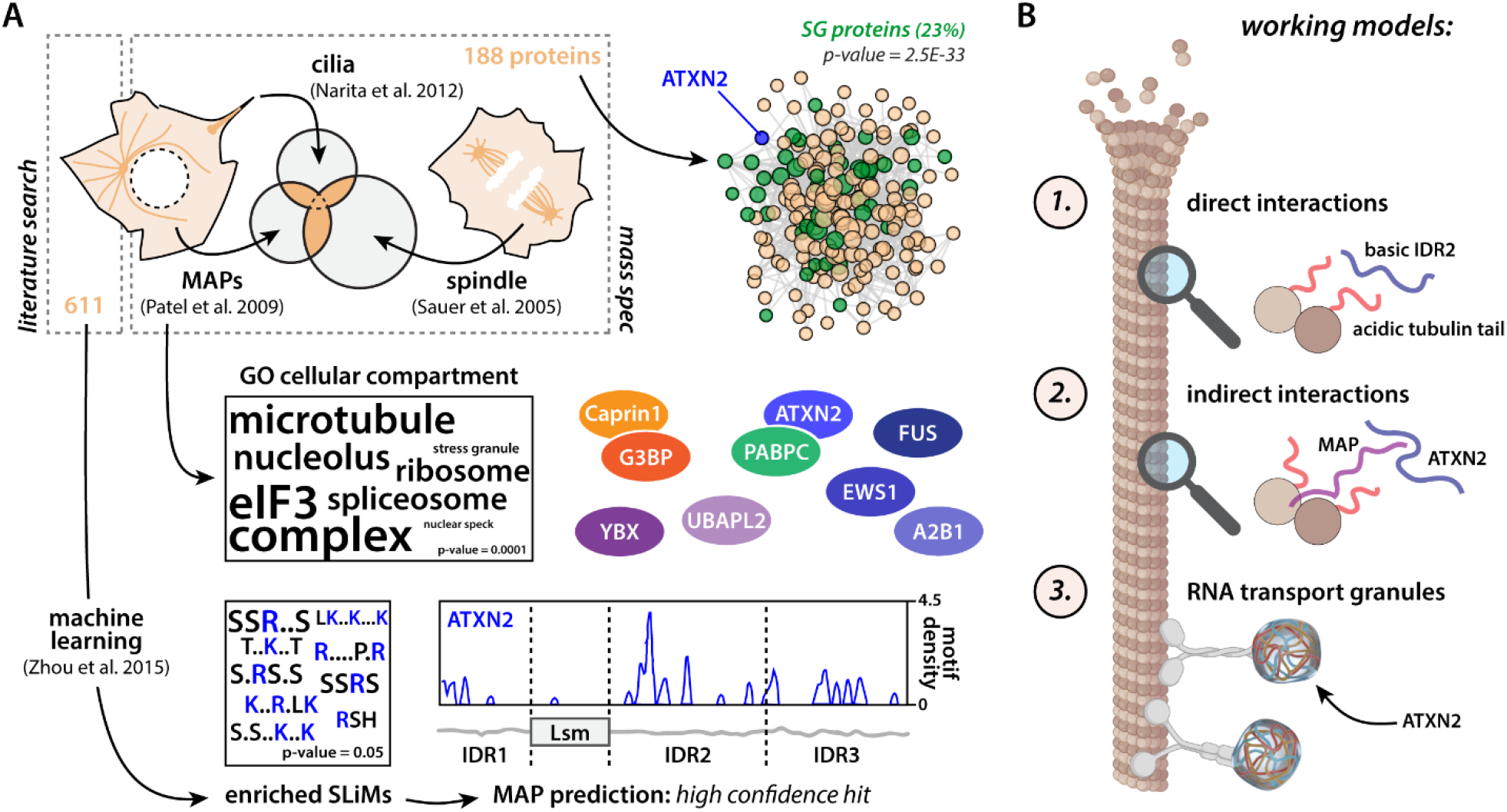
Possible interactions of stress granule proteins with the microtubule cytoskeleton. (A) Based on a literature search, Zhou et al. (2015) define 611 microtubule-associated proteins (MAPs), which are used to train a prediction algorithm based on enriched domains and short linear motifs (SLiMs)(Zhou et al., 2015). Word cloud shows enriched SLiMs with font size as a function of -log10(p-value). ATXN2 is a high confidence hit based on the enrichment of specific SLiMs, most pronounced in IDR2. Patel et al. (2009) identify a set of MAPs via mass spec (Patel et al., 2009). Performing GO enrichment on this dataset show an overrepresentation of terms associated with RNA granules and stress granules. Word cloud shows enriched terms with font size as a function of -log10(p-value). Several well-known stress granule proteins (colored ellipses) are found in this dataset. By overlapping the MAP dataset with mass spec datasets for spindle (Sauer et al., 2005) and primary cilia proteins (Narita et al., 2012), we find 188 proteins that were at least present in two out of three of these sets. This subgroup of proteins shares many physical interactions and shows a strong enrichment for stress granule proteins (Jain et al., 2016). Binomial test. (B) Possible interactions of ATXN2 with microtubules. (1) Direct interactions between acidic tubulin tails and the basic IDR2. (2) Indirect interactions of ATXN2 with disordered MAPs. (3) ATXN2 is found in RNA transport granules, which are tethered to the microtubules via molecular motors.

